# Independence of Visuomotor Functions Engaged in Visual Pursuit and Rapid Responses to Reach Errors

**DOI:** 10.64898/2026.02.28.708705

**Authors:** Renato Moraes, Jolande Fooken, J. Randall Flanagan

**Affiliations:** School of Physical Education and Sport of Ribeirão Preto, University of São Paulo, Ribeirão Preto, SP, Brazil; Department of Psychology and Centre for Neuroscience Studies, Queen’s University, Kingston, Ontario, Canada; Department of Psychology and Centre for Cognitive Science, Technical University Darmstadt, Darmstadt, Germany

**Keywords:** feedback gain, gaze, peripheral vision, reaching movements, sensorimotor integration

## Abstract

When reaching to a foveated target, peripheral vision of the hand can be used to make rapid, automatic adjustments to the ongoing reach movement, with the feedback gain being sensitive to features of the task and environment. These rapid corrective responses are also observed when gaze is directed to a stationary ‘gaze’ target located away from the reach target. In everyday contexts, reaching often occurs concurrently with other visual or visuomotor tasks, such as tracking a moving target. Yet it remains unclear whether engaging in such tasks affects the use of peripheral vision for hand guidance. Here, we compare rapid visuomotor corrective responses to visual perturbations during fixation and smooth pursuit, and test whether pursuit-related and reach-related visuomotor processes operate independently or compete for shared visual resources. Participants either fixated a stationary target or tracked a moving target while reaching toward a spatially dissociated reach target. During the reach, the visual representation of the hand was perturbed, requiring rapid corrective responses. We found that neither the onset nor the gain of reach corrections was modulated by gaze-task demands. Moreover, response gains were strongly correlated across tasks, indicating consistent individual response profiles that were independent of the gaze condition. Despite modest increases in position error and decreases in gain, participants largely sustained engagement with the visual tasks during target reaching. These findings demonstrate that smooth pursuit and reach-related visuomotor processing can operate in parallel without mutual interference, suggesting a functional independence between them.

**NEW & NOTEWORTHY:** In everyday life, reaching to an object can occur while the eyes are engaged in competing visual tasks. We show that engaging in smooth pursuit eye movements does not disrupt rapid visuomotor corrections during reaching. The onset and gain of corrective responses following perturbation were unchanged by gaze-task demands and were consistent across individuals. These findings demonstrate that pursuit and reach-related visuomotor processes can operate in parallel, supporting functional independence between these systems.

## INTRODUCTION

Visuomotor control in reaching involves the integration of visual information with motor commands to accurately move the hand toward a target location. To effectively use peripheral vision and gaze-related signals—including gaze proprioceptive signals—in reaching, individuals typically fixate the target throughout the reaching movement (1–3). These visual signals can be used to correct for movement errors following hand or target perturbation. Such reach corrections are characterized by a quick correction onset (80-150 ms) and flexible feedback gain, i.e., the magnitude of the corrective response (4). Whereas the onset of reach corrections occurs automatically and independent of features in the visual environment, feedback gains are sensitive to task context and goal (5–9).

In real-world situations, goal-directed reaching may occur in parallel with a competing visual or visuomotor task, such as identifying objects, monitoring environmental events, or tracking a moving object. Past research has shown that the visuomotor coordination, including corrective responses, remains intact when individuals fixate a ‘gaze target’ located away from the reach goal (10–12). However, it is unknown whether engaging in a more complex visual task, in which the eye position changes dynamically, influences our ability to use peripheral vision and gaze-related signals to direct the hand.

The aim of this study is to determine whether engaging in an ocular tracking task affects our ability to use gaze-related signals to direct and correct hand movements toward a peripheral reach target. We developed a task in which participants moved a cursor, displayed on a vertical monitor, from a start position to a reach target by moving the handle of a robotic manipulandum in a horizontal plane. In two conditions, participants either fixated a stationary gaze target (*fixation* condition) or tracked a moving gaze target (*pursuit* condition) prior to and during the reach movement. To assess the efficacy of the visuomotor control in reaching, we measured the onset and gain of the rapid visual feedback responses (mismatches between actual and predicted sensory feedback). Specifically, we included perturbation trials in which we jumped the position of the cursor to the left or right, while it passed beneath an occluder (3,13,14), and measured the gain of the resulting rapid corrective response using a force channel (9–11,15,16). The force channels provide a highly sensitive and reliable method for measuring corrective responses, as they are not compromised by limb dynamics (10,11,16).

Our paradigm allows us to address two alternative hypotheses. First, engaging in the ocular tracking task might interfere with the ability to use visuomotor signals to correct movement errors. Here, we would expect the timing and gain of the visuomotor corrective response to differ between the fixation and pursuit conditions. This would suggest that the two tasks operate dependently and thus visual resources cannot be fully used in parallel. Alternatively, performing the ocular tracking task might not interfere with the visuomotor control of reaching. Here, we would predict that individuals will be able to effectively integrate peripheral visual information and gaze-related signals to produce rapid, automatic corrective responses to visual perturbations, even while tracking a moving gaze target. Thus, the gain and onset of correction of these visuomotor feedback responses should be comparable in the pursuit and fixation conditions. This would suggest that ocular pursuit and reach-related visuomotor processing can operate in parallel without interference, indicating a degree of independence between these visuomotor functions.

## MATERIALS AND METHODS

### Participants

Twenty-four adults (21.7 ± 5.2 years, 16 women, 23 right-handed) participated in this study. Queen’s University students received one course credit, and community members outside the university received $15 for their participation. All participants had normal or corrected-to-normal vision, no upper-limb limitations, and no neurological conditions. The session lasted about one hour, and all participants provided written consent. The study was approved by the Queen’s University Research Ethics Committee. Data from four participants were excluded due to eye-tracker calibration issues and one due to possible nystagmus, resulting in a final sample of 19 participants (21.6 ± 5.2 years; 13 women; 18 right-handed).

### Apparatus

Participants operated a KINARM endpoint robotic manipulandum (BKIN Technologies, Kingston, ON, Canada) that allowed horizontal plane movements to control a cursor on a vertical plane monitor (70 × 39.5 cm; 1920 × 1080 resolution). The mapping between the robotic handle and the cursor resembled a standard computer mouse: moving the handle forward or right moved the cursor upward or right. Kinematic data and force data were sampled at 1,000 Hz. During the task, right eye movements were recorded using a monocular Eyelink 1000 system (SR Research Ltd., Kanata, ON, Canada) at 500 Hz.

### Visual stimuli

The hand position was represented on the screen as a circular cursor (1 cm in diameter) aligned with the handle of the robotic manipulandum. A 1-cm visual stimulus was equivalent to 1.5 visual degrees. Movements were made from a circular initial position (1 cm in diameter) to a circular target area (2 cm in diameter) located 25 cm above the initial start position (Fig. 1A). A 15 × 5 cm occluder was positioned between the start position and the target so that the far edge of the occluder was at the midpoint of the required reaching movement (i.e., 12.5 cm from the start position). Two visual targets were used to test how corrective responses depended on visual tasks: fixation (stationary gaze target) and smooth pursuit (moving gaze target). Participants had to maintain their gaze on the gaze target region throughout the trial so that the reaching movement was guided by peripheral vision (Fig. 1A). In the stationary gaze target, participants looked at a circular fixation target (0.5 cm diameter) (Fig. 1E). In the moving gaze target, participants were asked to track the displacement of the dot with their eyes as accurately as possible. The trajectory of the dot’s displacement was defined by equations described in previous studies (17,18). All trajectories had a period t of 6.3 s (fundamental frequency ω = 1 Hz; t = 2*pi/ω). The parameters used to generate the trajectories were replicated from a previous study (19).

**Figure 1.**
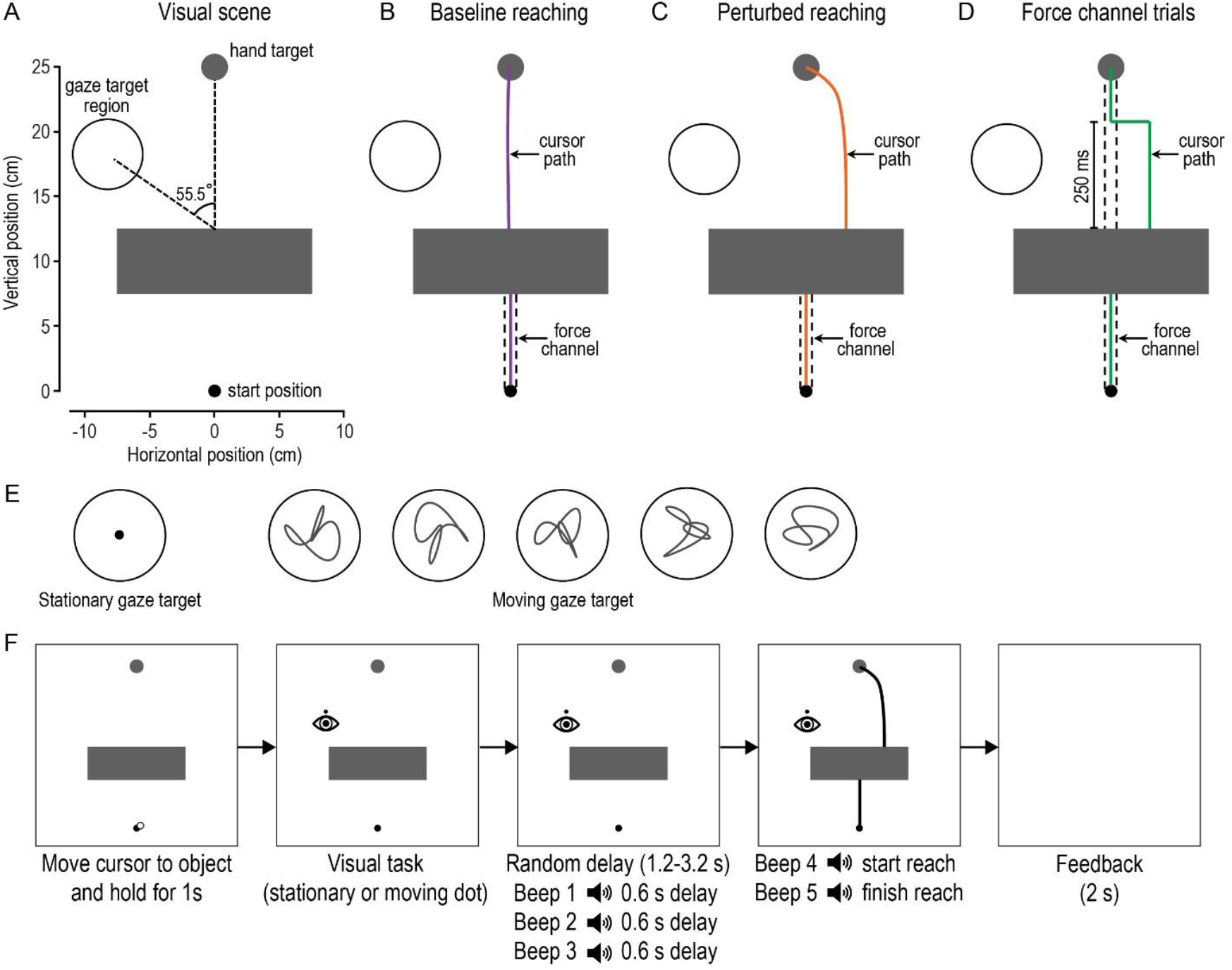
Experimental setup and conditions. (A) 2D view of the visual scene centred at the initial object location. For the visual task, participants either fixated a stationary circle (stationary gaze target) or tracked a dot that could move (moving gaze target) within the circle located in the left region (which was not shown to participants). The stationary gaze target was positioned at the centred coordinates x = -8 cm and y = 18 cm. Relative to the centre of the farthest edge of the occluder, this location placed the target 55.5° away at a distance of 9.7 cm. (B) Illustration of a trial without perturbation (baseline reaching). Participants performed the reaching movement from the starting position to the target while fixating or pursuing the gaze target with their eyes. (C) Illustration of a perturbation trial without a force channel following the perturbation (perturbed reaching). In these trials, the hand cursor was visually perturbed by shifting it 3 cm to the left or right after passing under the visual occluder. (D) Illustration of perturbation and force channel trial. In this condition, participants’ movements were constrained along a straight line from the start position to the target, which allowed the measurement of the forces applied to the virtual walls of the force channel (dashed lines). The cursor automatically returned to the straight line 250 ms after the perturbation onset in these trials. (E) Illustration of the stationary gaze target (left side) and the five dot trajectories used in the moving gaze target condition. (F) Temporal sequence of events in each trial. After the visual scene was presented, participants had to move the cursor to pick up the object. The visual task started 1 s after the object was ‘picked up’. The interval between beeps was fixed (0.6 s), and participants were instructed to begin to reach at the same time as beep 4 and finish the reach at the time of beep 5. After completing the reaching task, participants received feedback about the duration of the reaching movement.

### Procedure

Participants adjusted the seat height, positioned their chin and forehead on the support, and then completed the eye-tracker calibration. They first performed a familiarization block of 36 trials divided equally between the stationary and moving gaze targets, with only trajectory 1 (Fig. 1E) used for the moving target. Participants performed 12 trials of the non-perturbed reaching, 12 trials of the perturbed reaching, and 12 force channel trials in a pseudo-randomized order. For all participants, the first 5 trials involved non-perturbed reaching with a stationary gaze target, followed by 5 trials with a moving gaze target. The remaining conditions were fully randomized within the block. This sequence ensured gradual familiarization with all the steps involved in each trial.

#### General task

Once the visual scene appeared (object, target and occluder), participants moved the cursor to the initial position to pick up the object and held it there (Fig. 1F). After 1 s, the dot for the gaze target task appeared. After a delay between 1.2 and 3.2 s, five successive beeps (400 Hz; 80 ms) were presented 600 ms apart. The beeps served as a go cue: participants were instructed to start reaching on the fourth beep and arrive at the target on the fifth beep. On all trials, the cursor passed under the occluder. In perturbation trials, the cursor jumped 3 cm left or right below the occluder, requiring corrective adjustments when it reappeared to hit the target. The trial ended once the center of the cursor remained in the target for 300 ms. After each trial, a central message on the screen displayed movement time feedback (“good,” “very fast,” or “very slow”). Movement time from 1 cm above the initial position until 1 cm below the target was considered good between 400 and 700 ms. Participants were instructed to adjust their speed when feedback indicated it was not “good”. Trials outside this range were excluded (8.8%).

#### Non-perturbed reaching

In this condition, only the first 7.5 cm of movement was restricted by a mechanical channel (stiffness 2000 N/m, damping 0.2 N·m^-1^·s^-1^), after which the channel was slowed down for more than 50 ms (Fig. 1B). This restricted the initial reach, up to the near edge of the occluder, to a straight line and ensured that the cursor exited the occluder close to the line connecting the initial position and the target center (10,11).

#### Perturbed reaching

As in non-perturbed reaching, only the first 7.5 cm of movement was restricted by the mechanical channel (Fig. 1C). In this condition, the cursor jumped under the occluder and reappeared 3 cm to the right or left of the line between the start position and the hand target (10,11). Participants then had to correct the cursor’s trajectory to reach the hand target.

#### Force channel trials

We used force channel trials to assess the gain of corrective responses. In these trials, handle movement was restricted along a straight path from the initial position to the hand target position by a mechanical channel generated by the robot (Fig. 1D), allowing us to measure corrective forces exerted on the channel wall following visual perturbations. In these trials, the cursor exited the occluder either 3 cm to the left or right and was automatically shifted back to the straight path 250 ms after perturbation onset, consistent with previous work (9–11,15,16). Because this change occurred close to the time of correction, participants typically believed they were responsible for bringing the cursor back to the target. To avoid adaptive reductions in response magnitude, only 40% of trials used the force channel, randomly interspersed with non-channel trials.

Participants completed 15 trials for each of 10 experimental conditions: 2 perturbation directions (-3 cm [left] and +3 cm [right]) x 2 gaze tasks (stationary and moving) x 2 force channel conditions (without [perturbed reaching] and with [force channel trials]), totalling 8 conditions (120 trials). The two remaining conditions combined both gaze tasks with the non-perturb reaching, totalling 30 trials. All 150 trials were evenly distributed across 3 blocks in a single session, with trial order randomized within each block. Participants rested between blocks as needed to prevent fatigue.

### Data analysis

Eye and hand movement data were analyzed offline using custom MATLAB (version 2020b) routines. Hand movements were examined using the x and y positions of the robotic handle, filtered with a low-pass 3rd-order Butterworth filter (cut-off = 10 Hz). Eye movements were analyzed from calibrated screen-centred x and y coordinates, filtered with a 2nd-order low-pass Butterworth filter (cut-off = 15 Hz) and resampled to 1,000 Hz. Filtered data were used to identify fixations and gaze shifts.

The hand, dot, and gaze signals were differentiated to obtain velocity traces. Hand and gaze velocities were then low-pass filtered (10 and 25 Hz, respectively) to reduce noise from numerical differentiation. Hand onset was defined as the moment the vertical hand velocity first exceeded 5% of its peak value obtained during the reaching movement.

#### Force channel trials

To obtain a measure of the strength of corrections in response to cursor displacement (i.e., response gain), lateral forces in force channel trials were averaged over 180-230 ms after perturbation onset (9–11,16). For each participant and condition, the mean force following a leftward perturbation was subtracted from that following a rightward perturbation to obtain the corrective force difference (corrective force amplitude), computed separately for each gaze task. To calculate the onset times of force corrections, we compared individual force traces for left and right perturbations within each condition. Paired t-tests were applied at each time point after the perturbation onset to obtain the minimum p-value, then searched backward to locate the first time point with p < 0.001, which defined the onset of correction.

#### Visual task

Eye-position error was calculated as the root mean square error during fixation and smooth pursuit. This was obtained by subtracting the dot coordinates from the gaze coordinates. Visual gain between eye and gaze target velocities was computed for the moving gaze target task. Resultant velocities for both eye and gaze targets were calculated from their x- and y-velocity components. Eye velocity gain was calculated by dividing eye velocity (with catch-up saccades removed) by gaze target velocity.

Participants exhibited poor fixation or pursuit in some trials, characterized by large saccades that displaced the eyes significantly from the gaze target. Based on previous work (20), a 6-cm diameter boundary was defined around the center of the gaze target region (Fig. 2A, B). Trials with saccades landing outside this boundary (11.7% of trials) were excluded. Considering the two exclusion criteria (slow/fast movement time and large saccades), 19.1% of trials were removed from analysis.

**Figure 2.**
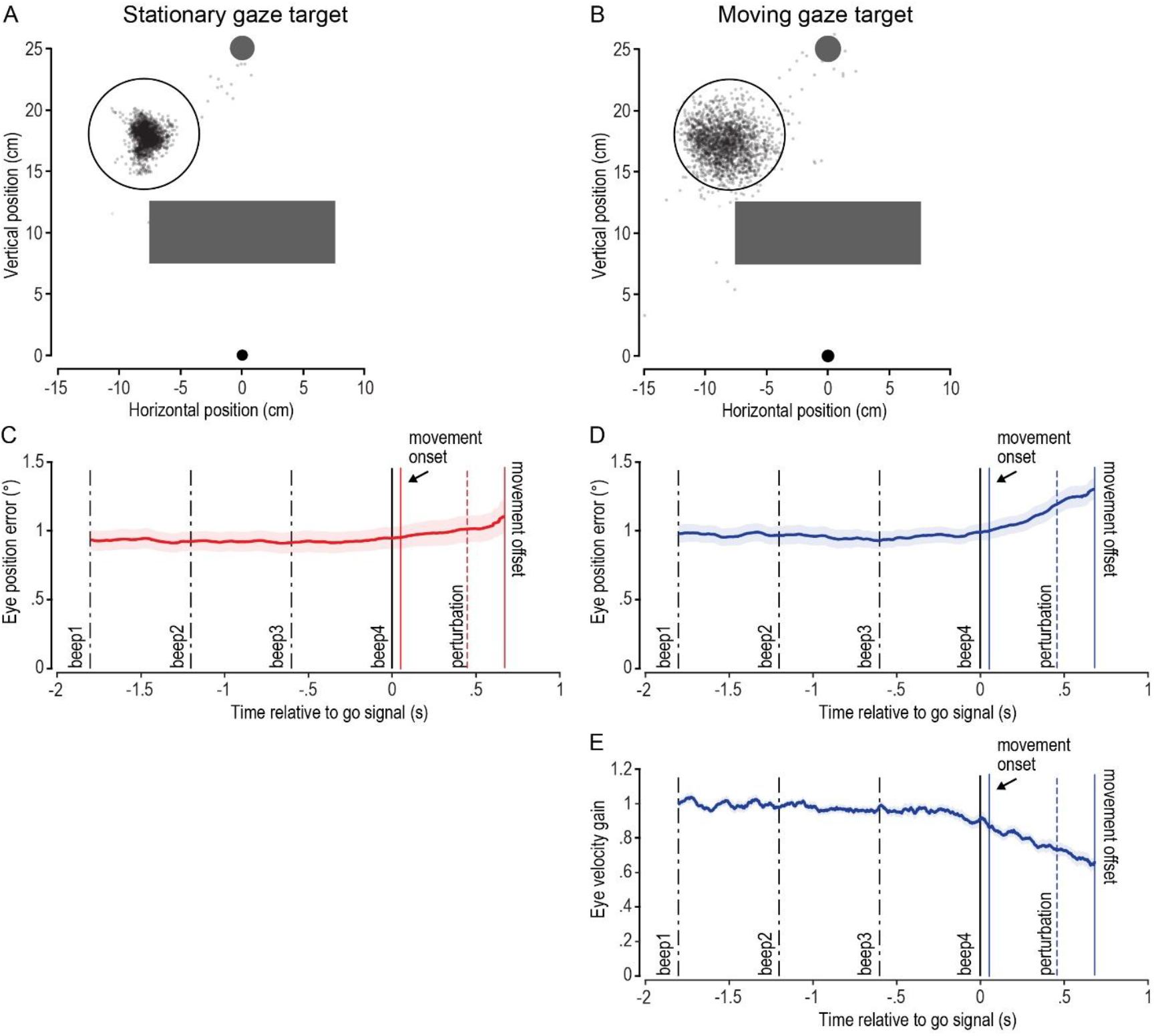
Eye position and saccade endpoints during reaching movements toward stationary and moving gaze targets. (A) In the stationary gaze target task, the saccade endpoints are shown in the upper panel as grey dots. Trials in which saccades landed more than 6 cm away from the gaze target zone (continuous circle on the left side) were excluded from analysis (see Methods). (B) The layout in the moving gaze target condition is similar to that in (A). The dark grey rectangle represents the occluder in (A) and (B). (C) and (D) Eye position error averaged across trials and participants for the stationary (C) and moving (D) gaze targets. (E) Eye velocity gain averaged across trials and participants for the moving gaze target. For (C), (D), and (E), the data are aligned to beep 4, and the shadow region around the mean corresponds to one standard error.

### Statistical analysis

Statistical analyses were performed in JASP (21). Statistical tests performed are described throughout the results section. In the text, results are presented as means (M) and standard errors (SE; M ± SE). The significance level was p < 0.05.

## RESULTS

### Participants can perform a secondary gaze task during reaching

Participants were instructed to fixate on the gaze target during stationary trials or pursue it during moving trials. Figure 2A and B show all saccade endpoints (grey dots) for stationary and moving conditions. Participants generally kept their gaze within the 6-cm boundary, though occasional large saccades occurred toward the hand target or elsewhere. Figure 2C and D show the average gaze position error across all trials and participants. During the period marked by the first four beeps, the average eye position error was about 1°. Following movement onset, it increased slightly by ∼0.1-0.2° for the stationary gaze target and ∼0.3° for the moving gaze target. For the moving gaze target, eye velocity gain was near 1 before beep 4 but declined gradually from movement onset to offset, reaching ∼0.7. Despite these shifts, the increases in position error and decreases in gain were modest, indicating that participants generally sustained engagement with the visual tasks.

### Hand movement kinematics are independent of gaze task demands

Hand kinematic parameters were similar between the stationary and moving gaze targets, indicating that the moving visual cue had minimal influence on the timing or kinematics of the reaching movement (Fig. 3). Fig. 3A shows cursor (i.e., hand) displacement in non-channel trials, where participants moved straight forward in both gaze conditions without perturbation. After left or right perturbations, trajectory adjustments appeared near the end of the reaching movement. The cumulative plots show data from the force channel trials (Fig. 3B-D). Kolmogorov-Smirnov tests revealed no significant difference between gaze conditions for any hand-movement parameter (vertical hand peak velocity: p = 0.609; movement duration: p = 0.559; movement onset relative to 4th beep: p = 0.688). On average, vertical hand peak velocity was 62.6 ± 0.2 cm/s, movement duration was 519 ± 2 ms, and reaching movement started 91 ± 7 ms after beep 4. Peak hand velocity occurred, on average, 12 ± 4 ms before the cursor left the occluder, showing that hand velocity was comparable across conditions around the cursor jump. Moreover, inter-individual differences in hand movement kinematics were consistent between gaze target conditions, as indicated by strong correlations for all measures (see the scatterplots shown in Fig. 3B-D; vertical hand peak velocity: r = 0.938; movement duration: r = 0.794; movement onset relative to 4th beep: r = 0.984; all p < 0.001).

**Figure 3.**
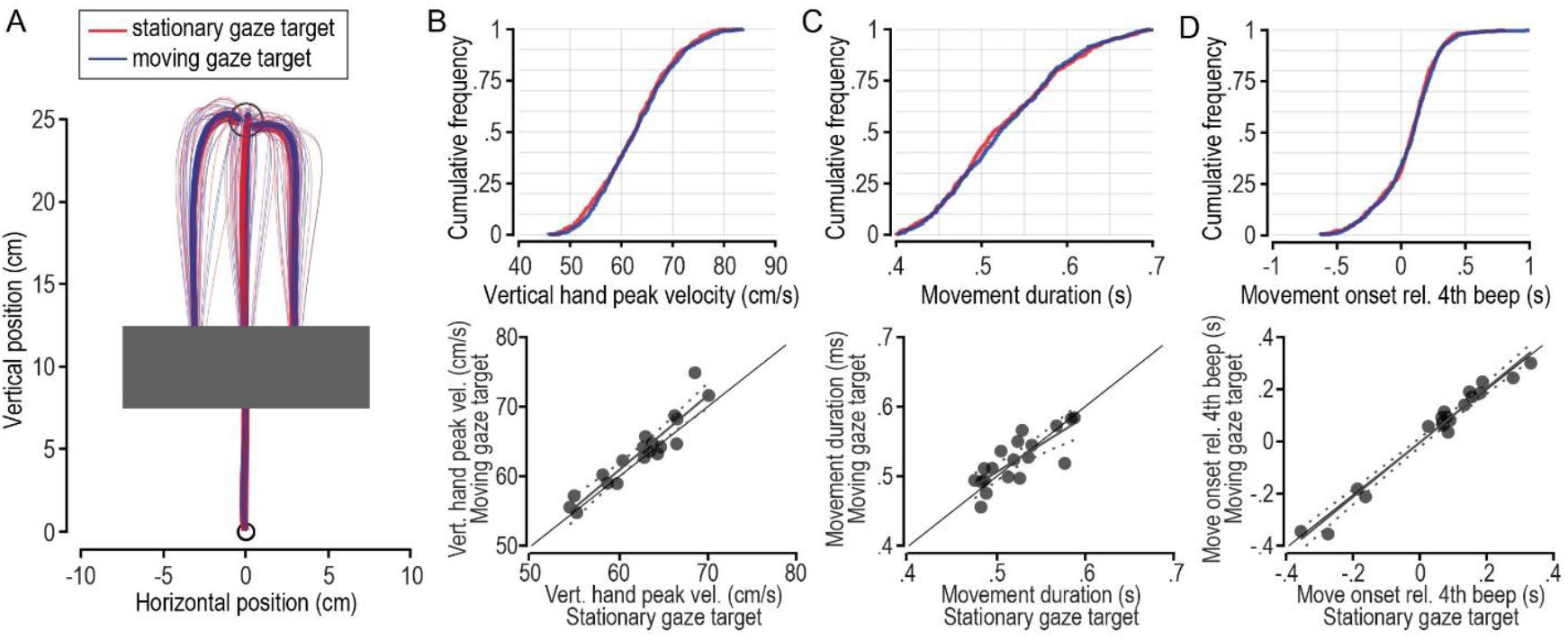
Hand kinematics and timing parameters under stationary and moving gaze targets. (A) Hand trajectories for each participant (thin lines) and average trajectory (thick lines) for the non-channel trials in the two gaze target tasks: stationary (red) and moving (blue). The grey shaded area indicates the occluder region. (B-D, top) Cumulative frequency distributions of vertical hand peak velocity, movement duration, and movement onset time relative to the fourth auditory beep (used as a temporal go signal), respectively, across participants and trials for the channel trials. (B-D, bottom) Scatterplots between stationary and moving gaze targets for the same variables described at the top. For these scatterplots, the median values of each participant and condition were used. Dashed lines indicate the 95% confidence interval for the line fitted to these data.

We also compared hand movement parameters between non-perturbation and perturbation conditions, using non-channel and non-perturbed reaching trials for the former (Fig. 3A). Kolmogorov–Smirnov tests again showed no significant differences (vertical hand peak velocity: p = 0.060; movement duration: p = 0.060; movement onset relative to the fourth beep: p = 0.524).

### Movement correction onset and gain are not modulated by gaze task demands

Fig. 4A-B shows, for the force channel trials, the forces exerted on the channel wall for the left and right perturbations in the two gaze targets for each participant (thin lines) and the averages across participants (thick lines). Fig. 4C shows the individual values of the correction onset times of the force responses during the reaching movement to the hand target. Correction onset times ranged from 140 ± 5 ms to 149 ± 4 ms for the stationary and moving gaze targets. The t-test for paired samples did not identify a significant difference between the gaze targets (t_18_ = -2.0, p = 0.057), indicating that the correction onset for the cursor perturbation was not delayed because of gaze target conditions. Additionally, we found no association between gaze target conditions for the correction onset (r = 0.399, p = 0.090), as illustrated in the scatterplot (Fig. 4C).

**Figure 4.**
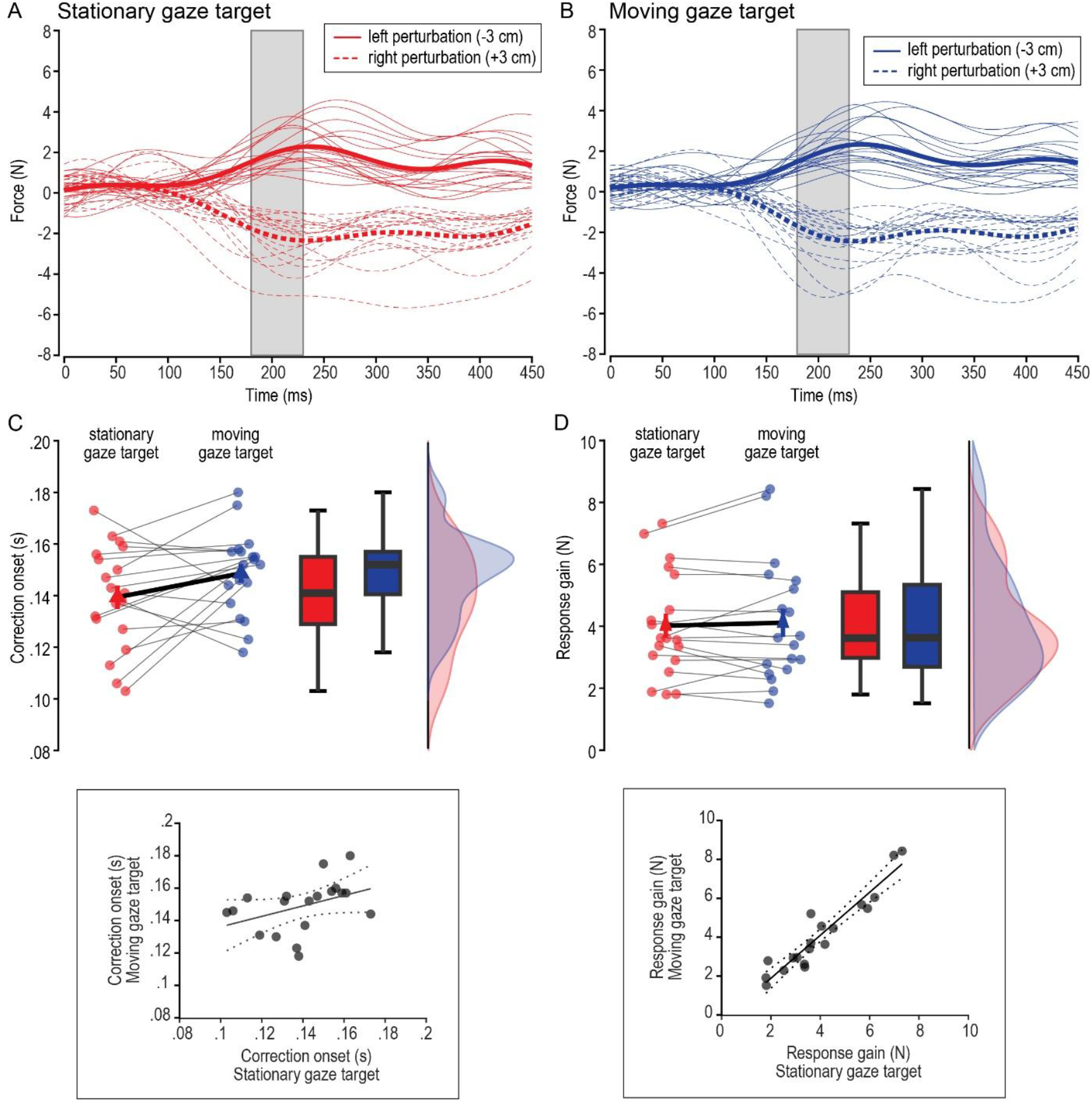
Visuomotor feedback gain and correction onset for stationary and moving gaze targets. Forces measured in force channel trials in the stationary (A) and moving gaze (B) targets. Responses are plotted for leftward (solid) and rightward (dashed) perturbation of the hand cursor during hand target reaching. Thin lines indicate the average forces for each participant, and thick lines indicate the average force across participants. The 0-ms corresponds to the instant of cursor jump. The grey area indicates the 180-230 ms interval used to average the force differences to obtain a single measure of response gain. The mean force values after a leftward cursor perturbation were subtracted from the mean forces after a rightward cursor perturbation to obtain the corrective force difference or response gain. The separation between left and right force time series was used to identify the onset time of correction following the cursor perturbation. (C-D, top) The top graph shows the correction onset (C) and the response gain (D) values for each participant (small dots) for both stationary (red symbols) and moving (blue symbols) gaze targets. The triangle indicates the mean, and the vertical bars indicate the standard error. On the right, the boxplots and probability density functions for both gaze targets are shown. (C-D, bottom) Scatterplots between stationary and moving gaze targets for the same variables described at the top. Dashed lines indicate the 95% confidence interval for the line fitted to these data.

As shown in Fig. 4 A-B, the interval used to calculate the mean force (180-230 ms) represents the period just after the onset of the response to the perturbation. The response gains were statistically compared using paired t-tests to examine the effect of the gaze target tasks on these gains. The results showed no difference in gaze target tasks for gain (t_18_ = -0.6, p = 0.545). Fig. 4D presents the mean values for each participant in the two gaze target tasks. The mean value was 4.02 ± 0.39 N for the stationary gaze target, and 4.12 ± 0.45 N for the moving gaze target. Unlike the correction onset, there was a strong association between response gain for stationary and moving gaze targets (r = 0.944, p < 0.001; Fig. 4D).

## DISCUSSION

The aim of the current study was to determine whether gaze-related signals involved in directing the hand movements toward a target would be disrupted by an active oculomotor tracking task. Participants either fixated a stationary target or tracked a moving target while reaching to a spatially dissociated reach target. During the reach, we perturbed the hand, requiring participants to make rapid corrections. We found that neither the onset nor the gain of reach corrections to cursor perturbations were modulated by gaze-task demands. Moreover, we found that response gains were strongly correlated across tasks, indicating consistent individual response profiles that were independent of the gaze-task manipulation. Together, these findings demonstrate that pursuit tracking and reach-related visuomotor processing can operate in parallel without interference, suggesting a degree of independence between these visuomotor functions.

### Gaze-related signals when reaching to visual targets

In the current study, participants successfully performed the secondary gaze task while reaching. When instructed to maintain fixation or pursue a moving target, they generally kept their gaze within the predefined boundary region, with only occasional large saccades away from the gaze target. Across conditions, eye position error remained low, approximately 1° during the preparatory period, and increased only modestly after movement onset. For the pursuit condition, eye velocity gain was near 1 before reach onset and gradually declined to about 0.7 by movement end. These small increases in position error and decreases in gain indicate that participants remained engaged and complied with gaze-task demands throughout the trials.

When naturally reaching to an object or visually cued location, humans typically fixate the target location throughout the reaching movement, a visuomotor behaviour that supports directing and guiding the hand towards the target (22). Previous work has shown that gaze-related signals—including proprioceptive signals from the eye and peripheral vision of the hand—support rapid, automatic corrections of movement errors when participants fixate on the reach target (2,3,14,23,24). Importantly, these rapid corrections also occur when participants fixate on a location dissociated from the current reach goal (11). Here, we show that corrections during goal-directed reaching are as fast and strong when participants—instead of fixating a stationary target—engage in an oculomotor tracking task, suggesting that continuous eye movements do not disrupt the use of gaze-related signals.

A classic problem in eye-hand coordination is determining which coordinate frame the visuomotor system uses to encode the spatial relationship between the sensorimotor apparatus and the external world (25). When fixating on the reach target, foveal vision of the target and peripheral vision of the hand can be used to directly compute a reach vector in visual coordinates. However, every time the eye moves, the relationship between the eye-centred and body-centred target location changes. Thus, computing the hand location relative to the target location in somatosensory coordinates requires the continuous integration of extraretinal signals for egocentric eye and hand position (26–29). The visual system also encodes allocentric, i.e., world-centred, target positions to support action across different task constraints and contexts, such as in memory-guided reaching (30–32). In our study, we find that engaging in an oculomotor tracking task does not impair the ability to rapidly correct reaching movements, suggesting that the integration of ego- and allocentric visual cues occurs automatically and early in the processing hierarchy, a process that may be similar to the continuous updating of visuospatial memory during smooth pursuit (33).

### Coordinated but independent control of eye and hand movements

Research on manual interception has shown that people naturally engage in smooth-pursuit eye movements when interacting with moving objects (34). When participants visually pursue an unpredictably moving target with gaze and are then instructed to reach towards and intercept the target, they not only continue to pursue the target after hand movement initiation but also tend to suppress catch-up saccades until the moment of interception (17). Moreover, engaging in smooth pursuit eye movements is linked to predictive motor preparation and correction of upcoming collisions (35,36). Taken together, these results highlight that visual feedback during smooth pursuit can continuously be integrated with online motor control.

Previous work has found that the control of eye and hand movements relies on shared early sensory and visuomotor neural processing (37–40). Behaviourally, it has been shown that attentional shifts to visually cued target locations can be allocated in parallel to upcoming eye and hand movements, even when the effectors are cued to spatially separate target locations (41–43). Thus, although the oculomotor and motor systems share early sensory input, motor plans and execution may operate in parallel (44–46).

### Gain put not correction onset are correlated within individuals

We found a within-participant correlation in feedback gain, but not in the onset of the correction, between the two visuomotor task conditions. These results are consistent with the idea that corrective feedback responses are initiated automatically—and as early as the processing hierarchy allows (47)— resulting in little variability across participants. At the same time, the gain of the response is known to be more sensitive to changes in visuomotor task demands (11,15,48) and thus it is not surprising it can also vary considerably across individuals. Similar to the feedback gain, we found that hand movement kinematics, that is, movement onset, duration, and peak velocity, were highly correlated within participants across tasks, suggesting that individuals have preferred movement signatures.

## Conclusion

Our findings demonstrate that smooth pursuit and reach-related visuomotor processing can operate in parallel without mutual interference, suggesting functional independence between these two visuomotor functions.

## DATA AVAILABILITY

Source data for this study are openly available at doi: 10.17605/OSF.IO/DFGQ2.

## ACKNOWLEDGMENTS

The authors thank Martin York for technical support.

## GRANTS

This work was supported by a Deutsche Forschungsgemeinschaft (DFG) Research Fellowship to JF (FO 1347/1-1), the Natural Science and Engineering Research Council of Canada (grant 156173 to JRF), the National Council for Scientific and Technological Development (CNPq, Brazil, grant 304056/2022-7 to RM), and the São Paulo Research Foundation (FAPESP, Brazil, grant 2019/21749-6 to RM).

## DISCLOSURES

The authors have no competing financial interests to disclose.

## AUTHOR CONTRIBUTIONS

RM, JF, and JRF conceived and designed research; RM performed experiments; RM, JF, and JRF analyzed data; RM, JF, and JRF interpreted results of experiments; RM, JF, and JRF prepared figures; RM and JF drafted manuscript; RM, JF, and JRF edited and revised manuscript; RM, JF, and JRF approved final version of manuscript.

## Notes

https://osf.io/dfgq2

### Competing Interest Statement

The authors have declared no competing interest.

